# Identical Dormancy Gene Mutations Reveal Unanticipated Relatedness Among Low-Chill Apples

**DOI:** 10.64898/2026.05.15.724974

**Authors:** Mujahid Hussain, Jugpreet Singh, Kevin M. Folta

## Abstract

Apples (*Malus* x *domestica*) are popular fruits grown in temperate regions of the world. The various genotypes must meet a specific threshold amount of cold exposure before they are competent to break dormancy, a quantity approximated as “chill hours”. Several varieties have been identified that exhibit an ultra-low-chill requirement, or more precisely shallow dormancy, breaking vegetative and floral buds early in spring in response to minimal cold exposure. These ultra-low-chill genotypes originated from the Bahamas (‘Dorsett Golden’,1960s), Israel (‘Anna’, 1950s) and Alabama, USA (‘Shell of Alabama’, 1880s). The separation in time and space implies that each would feature distinct genetic lesions that govern dormancy control, providing discrete mechanisms to incorporate a low-chill trait in variety improvement. However, analysis of microsatellites and ultimately genome sequence indicates that ‘Dorsett Golden’ and ‘Anna’ share strong concordance with the ‘Shell of Alabama’ genotype, as well as other ultra-low-chill varieties. Kinship analysis confirms that all are closely related, despite differences in year and place of origin. All three low-chill genotypes share common mutations in the *DORMANCY ASSOCIATED MADS-BOX1(DAM1)* gene, a known repressor of vegetative growth during dormancy. Genomic sequence diversity is observed among ‘Shell of Alabama’ individuals, including differences in *DAM1* that match differences in flowering time. The results of this study call into question the pedigrees of the ultra-low-chill apple germplasm and indicate variation in an otherwise narrow genetic base for use in future breeding efforts.

## Introduction

Apple (*Malus* × *domestica*, Borkh.) is a popular high-value fruit, grown in temperate areas of the world. Like most fruit tree crops in these regions generally require exposure to chilling temperatures to signal progression from dormancy to budbreak. But over the last century, several apple genotypes have been identified that flower and set fruit even in regions with minimal chilling temperature exposure. These varieties hold special importance in tropical and sub-tropical regions because they possess the genetics to release dormancy and produce early flowers and fruits with harvests preceding intense summer heat. These accessions may prove valuable in breeding programs that seek to create varieties for the tropics and sub-tropics, as well as provide traits that may be useful into new temperate varieties as winters warm from climate change (Hussain et al., in review).

Seasonal dormancy is typically defined by two phases – endodormancy and ecodormancy. Next year’s buds develop throughout the end of summer, as days get shorter and cooler, ultimately ending in in dormancy to traverse winter months. Endodormancy is the state of growth arrest where dormant buds accumulate “chill hours”, roughly defined as time spent between 0° and 7°C. After a threshold of hours is satisfied, the buds enter ecodormancy and are competent to respond to “heat units”, time accumulated under warm temperatures. Together these mechanisms position budbreak in favorable springtime conditions, in sync with other varieties and seasonal pollinators. Apple varieties vary in their endodormancy chill requirement from under 100 hours to temperate varieties requiring well over 1000 (González Noguer et al, 2023).

There is a long history of low-chill apple cultivation in places like Isreal (Holland et al, 2015). Decades to centuries later several apple varieties are described in catalogs and websites as “low chill” that exhibit extremely shallow dormancy (referred herein as “ultra-low-chill”; ULC). These originate from the 1880s to 1960s, from notably disparate locations. The main varieties include ‘Anna’, ‘Ein Shemer’, ‘Dorsett Golden’ and ‘Shell of Alabama’. These varieties, along with several other regional niche varieties, are recommended as sympatric plantings for efficient pollination, as they all flower in unison. The cultivars ‘Anna’ and ‘Ein Shemer’ were introduced in Israel by breeder Abba Stein in 1959, who sought to create an apple that would flower in a generally hot, Mediterranean desert climate (Holland et al, 1996) rarely exceeding 300-400 chill hours. The popular story is that the variety “Dorsett Golden” was identified in the Bahamas by Mrs. Irene Dorsett in the 1950s. This chance seedling from ‘Golden Delicious’ thrived and set fruit in the island heat, and in the early 1960s was imported to Florida where it was cultivated commercially and studied for breeding potential (Miller & Sherman, 1980). Another ULC apple is the ‘Shell of Alabama’, introduced in the 1880s in southern Alabama, just over the Florida panhandle state line. These “Shell Apples” were grown by Mr. Green Shell, an orchardist who daily sent two loads of apples in barrels to Brewton, AL for distribution northward in June and July (Calhoun, 2010). Shell Apples were frequently cultivated in the American South in the late 1800s.

Warmer winters and an increasing interest in diversifying tree crops in tropical and sub-tropical areas have refocused attention on the potential for ULC apple improvement. The primary hypothesis is that the low chill trait was selected independently, as every low chill cultivar has a history of discovery and cultivation in their respective regions. The separation of time and space suggests that the trait would likely be governed by different mutations and possibly separate genes, and may relate to other varieties with limited cultivation known to flower after less cold exposure.

To answer this question, we examined the genetic relatedness between the well-known ‘low chill’ varieties and other southern USA apple genotypes that have been described as ULC. Candidate genetic mechanisms that underlie the ULC trait were also explored. The results indicate that the three main varieties, presumed to be significantly different based on separation in timing and location of origin, are actually highly related, suggesting it is necessary to redraw pedigrees and correct the legends of low-chill apple origins. Establishing these genotypic relationships and delineating common/distinct ‘low chill’ mechanism(s) may aid future apple breeders in parental selection, and open opportunities into the specifics of molecular mechanisms of dormancy control.

## Materials and Methods

### Plant materials

The study examined 36 genotypes chosen based on their chill hour requirement (Table. I) that were grouped as ultra-low chill (100-200 chill hours), low chill (200-300 chill hours), moderate chill (300-500 chill hours), and high chill (> 500 chill hours). Trees were obtained from either commercial nurseries or were grown from grafted budwood and maintained on a farm in Archer, FL on either Geneva 890 or EMLA111 rootstocks. Leaf material of ‘Shell of Alabama’ was obtained from independent sources as outlined in Table II.

### Simple Sequence Repeat Analysis

Fresh leaves were collected for DNA extraction with the method described by Doyle & Doyle (1987) with modifications. To determine the initial relatedness of all genotypes, SSR analysis was performed using a set of fluorescence-tagged primers known to detect robust microsatellite polymorphisms in apple (Supplementary Table I). Amplicons were sent to the University of Missouri Genomics Technology Core (https://mugenomicscore.missouri.edu/) for Fragment Analysis using the ABI 3730xl DNA Analyzer. Peak data were analyzed to determine the fragment sizes in base pairs (bp) using a Python script. The two highest Peaks (biallelic) were filtered to remove the background noise based on peak size and height. Peaks with expected size range were included for analysis. A minimum peak height threshold was set to exclude spurious low intensity fluorescence signals. To determine the relatedness, the distance matrices were calculated based on alleles size calls among the various genotypes using custom Python (3.8) script.

### Relatedness based on whole genome sequencing

To maximize nuclear DNA sequence coverage, nuclei were isolated from fresh leaves of ‘Dorsett Golden’ and ‘Shell of Alabama’ using the method described by Li et al (2020) with the modifications shown in Supplementary Data. For library preparation and sequencing, the samples were sent to NextGen DNA Sequencing core at Interdisciplinary Center for Biotechnology Research (UF | ICBR) at the University of Florida, Gainesville, Florida (https://biotech.ufl.edu/next-gen-dna/). The libraries were prepared using NEBNext® Ultra™ II DNA Library pre kit following the manufacturer’s instructions (E7645S; New England Biolabs, Inc., USA) Genomic DNA was sequenced using Illumina NovaSeqx platform. For comparison, with moderate and high chill cultivars, the raw reads of ‘Anna’ (SRR3538832), ‘Gala’ (SRR10983009), ‘Red Delicious’ (SRR23461140), ‘Antonovka’ (SRR24967679), and ‘Honeycrisp’ (SRR17323561) were retrieved from NCBI SRA (https://www.ncbi.nlm.nih.gov/sra).

After necessary trimming, filtering, removal of adaptor sequences, and quality check using default parameters of trim galore (v0.6.10), fastQC (v0.12.1), and multiQC (v1.33) Linux packages, the reads were aligned to the reference ‘Golden Delicious’ (GDDH13 v1.1 genome assembly) apple genome, retrieved from Phytozome (https://phytozome-next.jgi.doe.gov/info/Mdomestica_v1_1). The Burrows-Wheeler Alignment (BWA; v0.7.17) tool (Guo & Huo, 2024) was used for alignments with default parameters. The alignment files were used for variant detection using the bcftools (v1.22). Briefly, the aligned reads were processed for genotype likelihood using bcftools mpileup which were used for variant calling using bcftools call using -m option for multiallelic variant calling with diploidy (--ploidy 2) setting assuming all cultivars are diploid (Lefouili & Nam, 2022).The resulting files generated were used for downstream comparative analysis.

To determine the genomic relatedness between the apple genotypes, a high-quality SNP dataset was used to conduct an identity-by-descent (IBD) analysis in PLINK v1.9 using -- genome parameter. The sharing probabilities of zero (Z0), one (Z1), or two (Z2) alleles identical by descent and relatedness coefficient (PI_HAT) were estimated. Furthermore, to estimate genome-wide similarity, a distance matrix was created using DST (distance based on genetic similarity) parameter in PLINK (v1.9) and a heatmap was generated using custom Python (v3.8) scripts (Figure. 2).

### Identification of identical mutations among ultra-low-chill cultivars

The variants shared among all the low chill genotype sequences were filtered out using the isec function of bcftools. (v1.22) These variants were unique to ‘low chill’ germplasm and were absent in the moderate to high chill apple cultivars. These variants were annotated using SnpEff (https://pcingola.github.io/SnpEff/) and an GDDH13 v1.1 genome annotation database built with gffread (https://github.com/gpertea/gffread) package (v0.12.7). The variants were annotated to define major effect mutations including gain and loss of stop codon, frameshift, gain and loss of translational start, developing an errant splice donor site, missense mutation, and splice acceptor variants.

The selection of candidate genes was made by filtering out variants that are not linked to dormancy, bud break, and flowering time pathways. First, the annotated shared variants of ULC genotypes were analyzed with InterPro Scan (https://www.ebi.ac.uk/interpro/) for identification of domains associated with uncharacterized gene IDs. To predict the functions of these genes, the *Arabidopsis thaliana* and peach (*Prunus persica*) orthologs were identified using the EnsemblPlants (https://plants.ensembl.org) database. Genes associated with dormancy, bud break, and chill hour requirement were identified based on InterPro domains and descriptions of *A. thaliana*, and *P. persica* orthologs. Broad regulatory proteins were excluded to narrow down the parameters of candidate gene identification.

### Development of a ULC-Associated Marker

For molecular assisted breeding, we developed a marker to quickly identify that either seedling has low chill genotype or not. We identified indel downstream of *DAM1* SNPs which co-segregate with *DAM1* mutations. We analyzed the loci of *DAM1 and* surrounding indels using our annotated variants using custom Python script. The Pairwise linkage disequilibrium (LD) among SNPs and indel was also calculated using --geno-r^2^ and --min-r^2^ 0.0 in VCFtools (v0.1.16).

To test the association between the deletion and ULC cultivars, DNA was extracted from ULC ‘Anna’, ‘Dorsett Golden’, ‘Shell of Alabama’ (different accessions; see Table II), ‘Ein Shemer’, and ‘Vered’ as well as the low chill cultivars ‘Mutsu’, ‘Cripps Pink’, and ‘Yellow Bellflower’ and high chill cultivar ‘Golden Delicious’ using method described in supplementary file. The identified indel was amplified using forward (5’GATTAAGCCAGGCCGGTAG3’) and reverse (5’GAGCACATTGTGAGGCTTAATG3’) primers. The same primers were used to track inheritance of the indel and associated mutations in a segregating set of F_1_ seedings from a cross between ‘Anna’ and ‘Reverend Morgan’.

For the validation of co-segregation of both *DAM1* SNPs and the identified indel, DNA sequences were obtained from the colonies of subset of ULC, low chill, and F_1_ cross of ‘Anna’ and ‘Reverend Morgan’. The SNP-containing region was amplified using forward (5’GACAAGATGTCGGAGCAATCA3’) and reverse (5’GGTAAGCAACTACCACCAAGAA3’) primers, and the amplicons were ligated into TA TOPO T/A cloning vectors (Invitrogen) following the manufacturer’s protocols, and transformed into *E. coli* for amplification. The plasmids were isolated using a Qiagen miniprep kit and was sent for Sanger Sequencing to GENEWIZ (www.genewiz.com).

## Results

### Genetic relationships between low chill apples using SSR markers

Microsatellite variation was first used to assess the relatedness between ULC genotypes, as well as compare them to other apple cultivars, including varieties described as “low chill” and other southern USA varieties. A set of fluorescence tagged primers was used to amplify well-established polymorphic Simple Sequence Repeats (SSRs) in apple (Table I). The results show that ‘Anna’, ‘Shell of Alabama’, ‘Dorsett Golden’, ‘Elsa Sweet’, and ‘Joy’s Apple’ share high allelic similarity with genetic distance from 0.92 -0.93 (Figure 1), whereas other southern USA low chill materials do not. ‘Carolina Red June’ is presented as a comparator, along with ‘Chestnut Crab’, a Minnesota variety of crabapple, representing a presumably distant genotype. The allele-based similarity and pairwise distance revealed high similarity among ULC genotypes (Supplementary Figure 1), defining two distinct groups. (Supplementary Figure 1).

**Figure 1.**
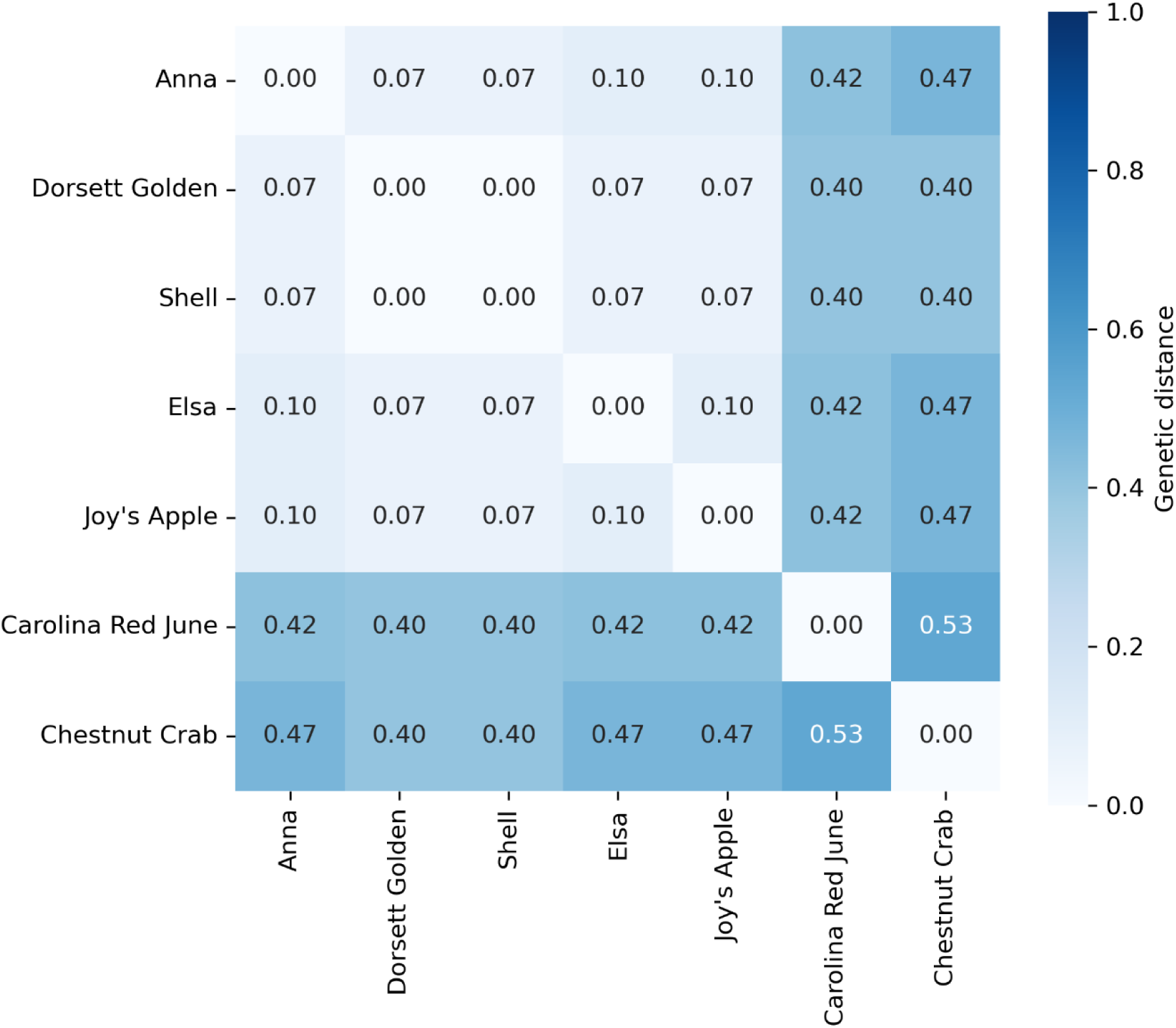
Heatmap showing the genetic distance among Shell-like low cultivars ‘Anna’, ‘Dorsett Golden’, ‘Joy’s apple’, ‘Elsa Sweet’, ‘Shell of Alabama’, compared to a low-chill southern variety ‘Carolina Red June’ and a genetically distant ‘Chestnut Crab’, based on SSRs allelic data. The colors and the numbers on the grids represent the genetic distances, and closer numbers among cultivars show high relatedness.

**Figure 2.**
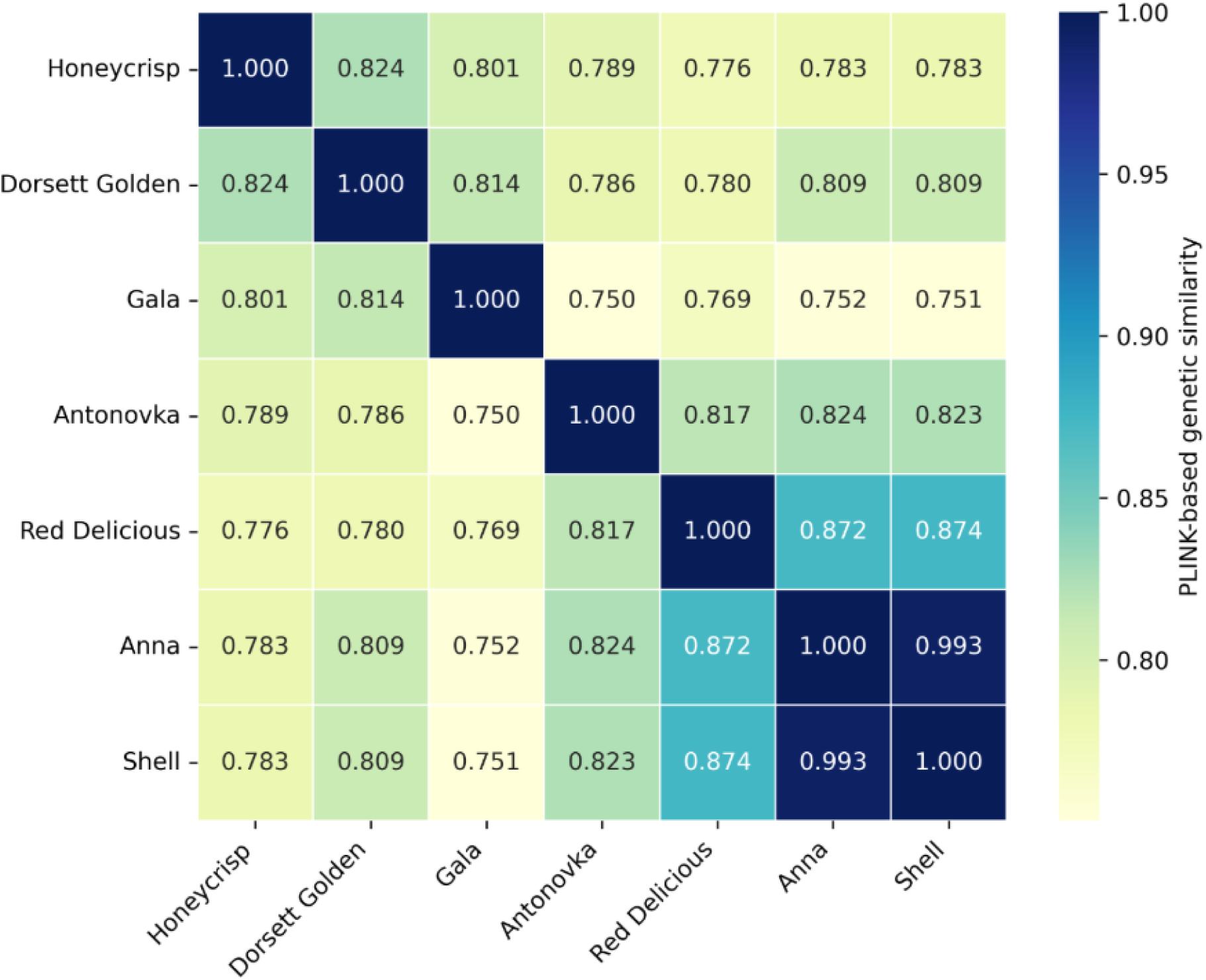
Heatmap depicting genome-wide similarity based on whole genome variant data. The pairwise genetic distances were calculated based on PLINK. The numbers in the grid depict similarity among pairs of cultivars ranging from 0-1, with higher number means higher relatedness.

**Table I.**
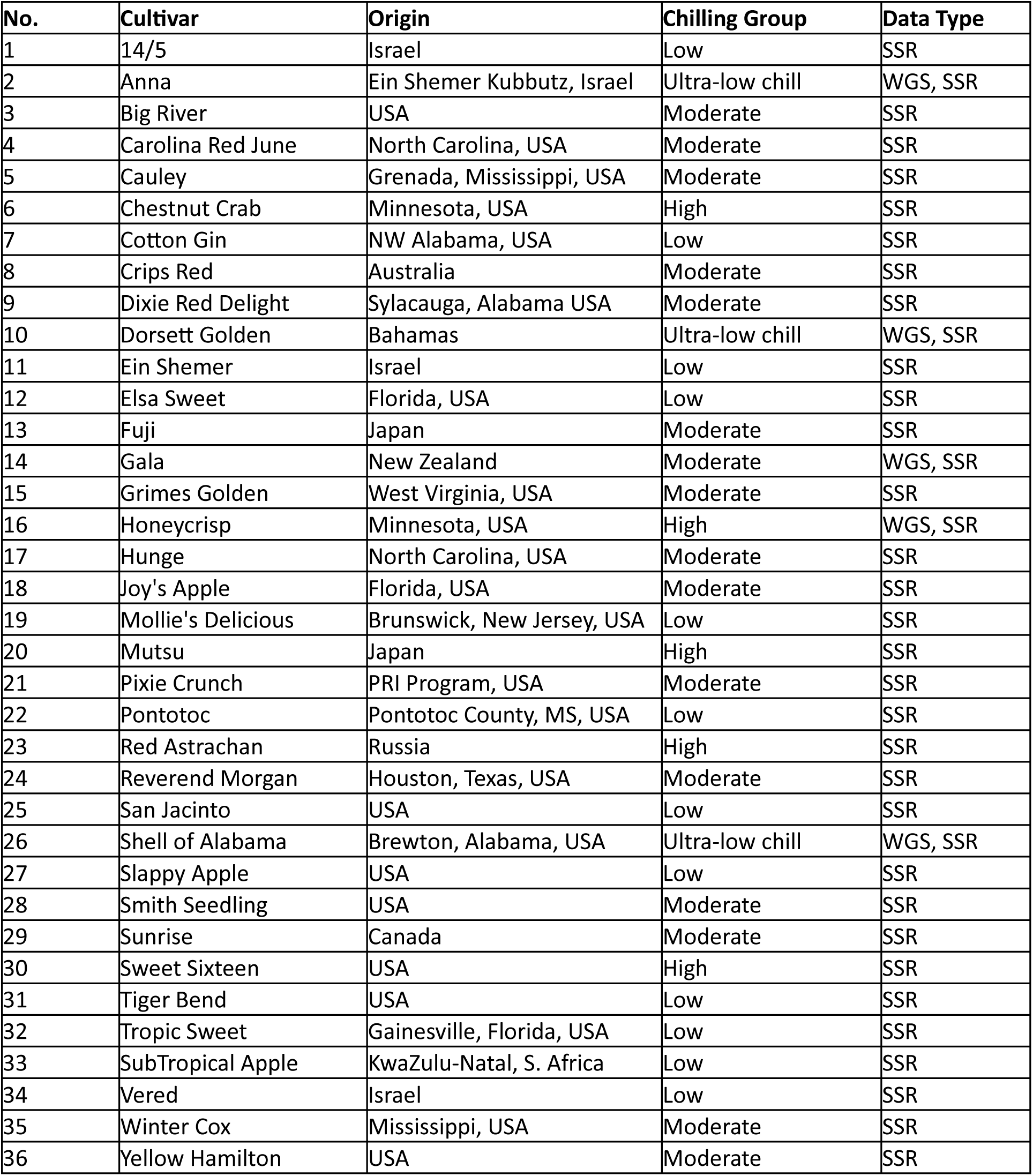
Apple cultivars used in whole genome sequencing and SSR analyses.

### Low chill cultivar relatedness using whole-genome variant analysis

The SSR result clearly indicated strong genetic relatedness between the ULC genotypes. However, limited number of SSRs may cause bias in calculations of genetic relationships. To allow for a higher resolution comparison between these varieties, a whole genome sequencing approach was employed. The genomes of ‘Dorsett Golden’ and ‘Shell of Alabama’ were compared against publicly accessible genomic data from ‘Anna’, ‘Gala’, ‘Red Delicious’, ‘Honeycrisp’, and ‘Antonovka’. Identity-by-descent (IBD) analysis revealed remarkable similarity among ULC genotypes. ‘Anna’ and ‘Shell of Alabama’ showed highest genetic similarity with DST ≈ 0.99 and PI_HAT ≈ 0.98 potentially indicating a clonal or bud-sport relationship (Figure 1; Supplementary Table II). However, ‘Dorsett Golden’ showed a moderate similarity with ‘Anna’ and ‘Shell of Alabama’ with DST ≈ 0.81; PI_HAT ≈ 0.57. The two other cultivars ‘Red Delicious’ and ‘Antonovka’ also shared moderate similarity with ‘Anna’ and ‘Shell of Alabama’ with DST 0.82 and 0.87, respectively.

Remaining cultivars displayed more divergence from the ULC cultivars. For example, ‘Gala’ showed ≈ 0.75 similarity score with PI_HAT ≈ 0.50 with ‘Anna’ and ‘Shell of Alabama’. Similarly, a distant relationship was found among pairs of ‘Dorsett Golden’ and

‘Red Delicious (similarity ≈ 0.77), ‘Antonovka’ and

‘Honeycrisp’ (similarity ≈ 0.78), and ‘Shell of Alabama’ and

‘Honeycrisp’ (similarity ≈ 0.78; Figure 1). Based on IBD analysis, the genotypes studied formed three groups, the seemingly clonal relationship between ‘Anna’ and ‘Shell of Alabama’, the moderately related group in case of ‘Dorsett Golden’ and ‘Red Delicious’, and third group showing genetically distant background like ‘Gala’ and ‘Honeycrisp’.

### A Common Mutation in a Known Dormancy Control Gene

Members of the *DORMANCY RELATED MADS BOX* (*DAM*) genes have been shown to repress dormancy in a variety of rosaceous tree crops, including peach (Bielenberg *et al*, 2008), pear (Boldizsár *et al*, 2020), and apple (Moser *et al*, 2020). The work by Moser and colleagues (2020) demonstrated using transgenic gain- and loss-of-function approaches that the *MdDAM1* gene has a major influence on dormancy control. Therefore, scrutiny of this gene was warranted. Analysis of ‘Anna’ and ‘Dorsett Golden’ gene sequences show matching mutations in *MdDAM1* (MD15G1384500) not found in modern and higher-chill cultivars, including ‘Gala’, ‘Red Delicious’, ‘Antonovka’, and ‘Honeycrisp’ (Figure 3). The SNPs at the 1st (position: 47466812) and 7th (position: 47475865) exons create amino acid changes that may be contributing to the dominant phenotype. ‘Shell of Alabama’ flowers in concert with ‘Anna’ and ‘Dorsett Golden’, and contains the identical mutations (Figure 3).

**Figure 3.**
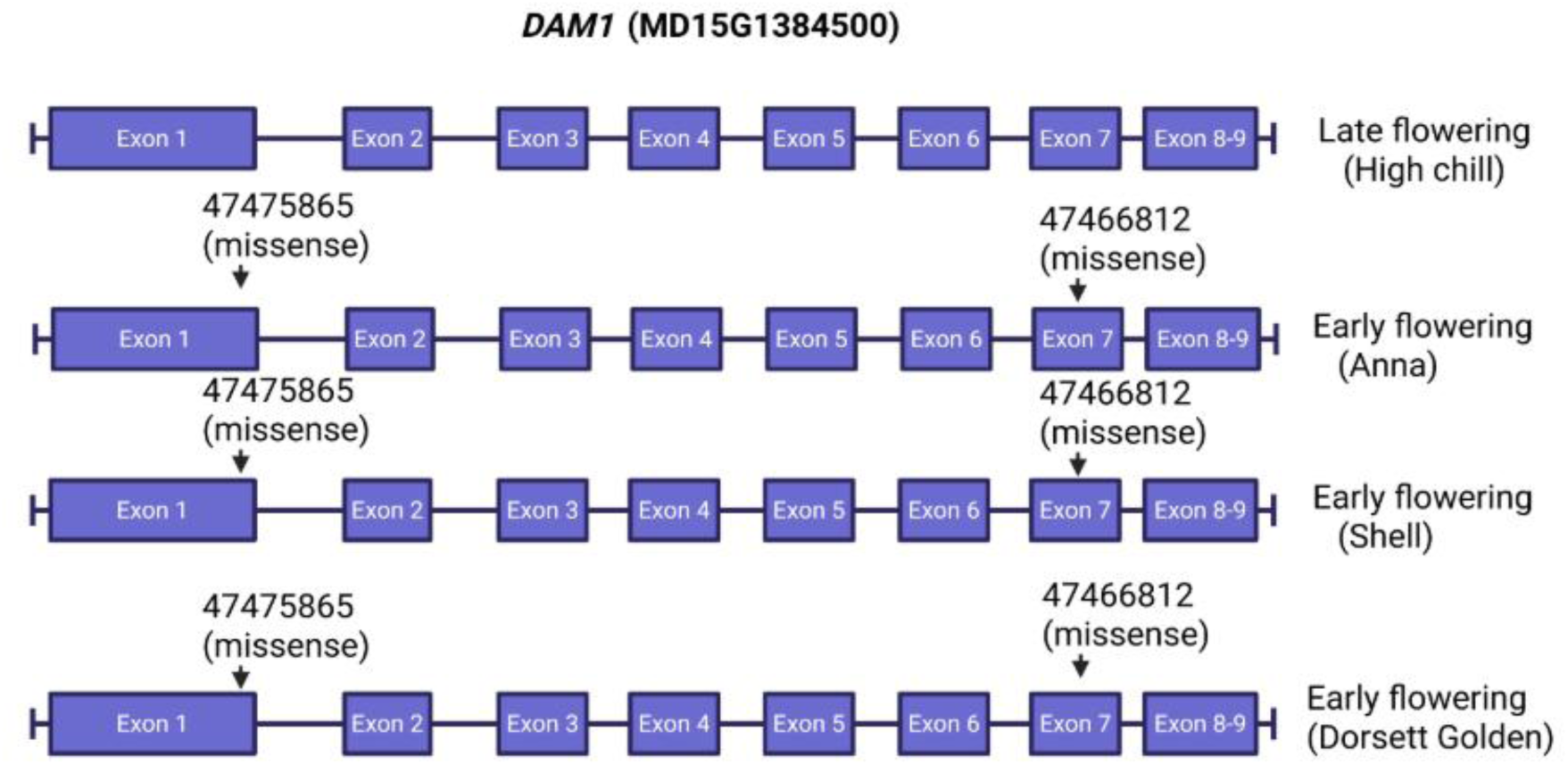
The mutations in *DAM1* gene. The non-ultra-low-chill varieties such as ‘Gala’, ‘Honeycrisp’ and ‘Antonovka’ present a consistent sequence in *DAM1*. However, the ultra-low-chill varieties ‘Anna’, ‘Dorsett Golden’, and ‘Shell of Alabama’ (Shell) show SNPs at exon1 and exon 7 that affect amino acid composition in key interaction domains.

### A Molecular Marker for ULC

The SNPs may prove useful in the breeding of ULC apples, as ULC is a dominant trait. Genetic improvement may be rapid because at least half of the individuals in the first generation will maintain the ULC trait. However, SNP detection requires special techniques and/or equipment, and a simple PCR based method would hasten detection of likely ULC genotypes. Analysis of the genomic regions adjacent to the *MdMAD1* gene revealed a 36 bp indel 18,204 bp from the downstream paralogue of *DAM1* SNP in *dam1* (Figure 4A). To test if the indel was present in other ULC genotypes, and if the marker segregated with the *MdDAM1* mutation, PCR primers were designed flanking the region, detecting either a 187 bp fragment in genotypes containing the *dam1* mutations and the wild-type 223 bp fragment alone in non-ULC material. The pairwise LD among SNPs and indel also shows that these are perfectly co-segregating (Table IV). The approach was tested against 22 genotypes; a sample of 10 is shown in Figure 4B. The 187 bp fragment containing the deletion is detected in ULC genotypes such as ‘Ein Shemer’ and ‘Elsa Sweet’ and not in higher chill materials. All ULC genotypes were heterozygous for the mutation. The inheritance was tested in progeny from an ‘Reverend Morgan’ x ‘Anna’ cross. Seedling DNA was screened for the 36 bp indel, ‘Anna’ being heterozygous for the marker, while ‘Reverend Morgan’, a highly heat tolerant low-chill seedling of ‘Granny Smith’, does not contain the deletion or the corresponding *dam1* mutations. The results show that the lower molecular weight band was detected in 6 of 10 seedlings, consistent with Mendelian inheritance.

**Figure 4.**
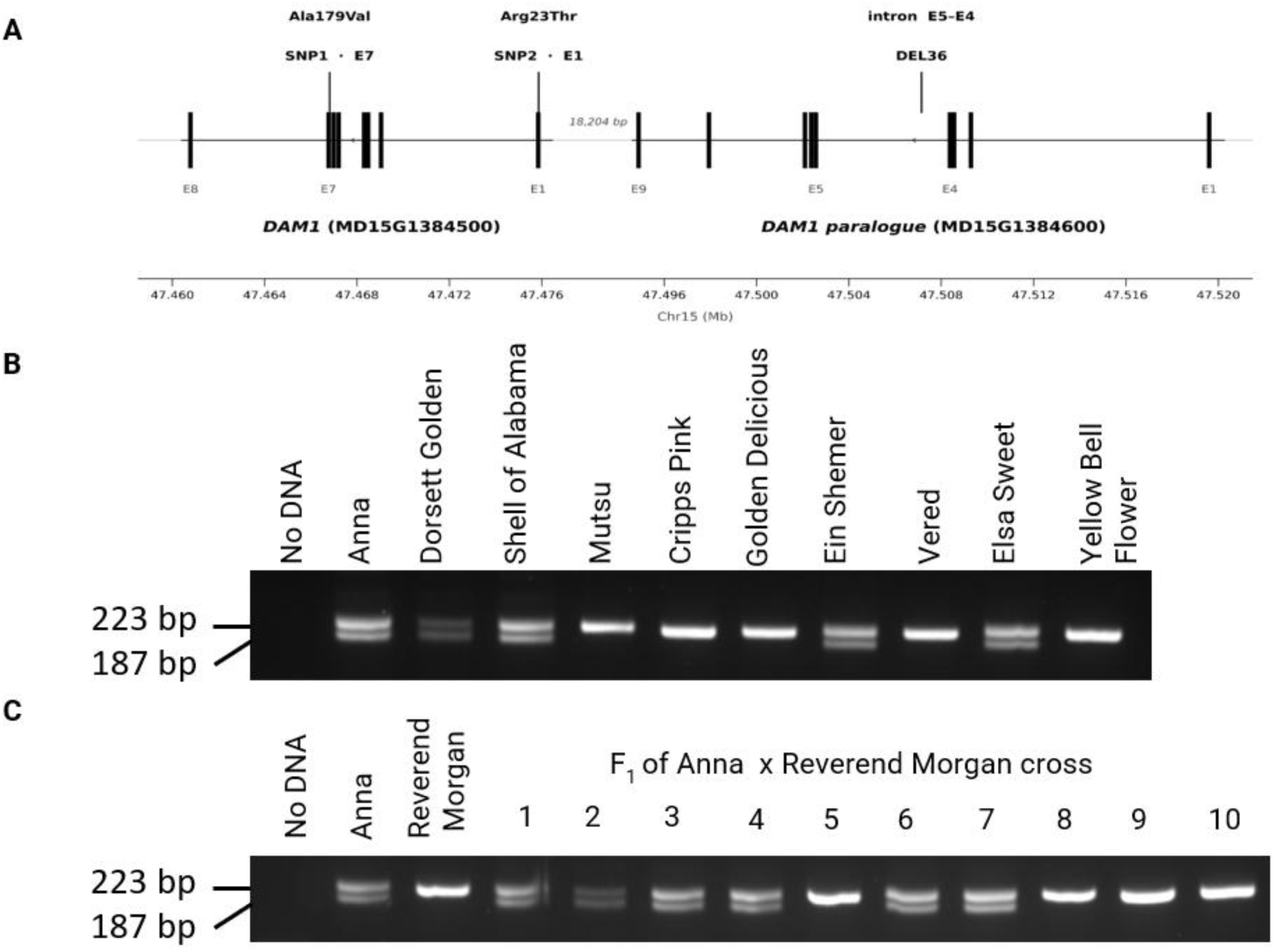
Identification of a PCR-detectable indel that segregates with the ultra-low-chill allele. Panel A shows the genomic context of a 36 bp deletion present 18,204 bp downstream of the *dam1* SNP on Chromosome 15. Panel B shows its presence/absence in a series of ultra-low-chill and other apple genotypes, the wild-type product represented by a 223 bp product, the 187 bp band co-segregates with the *dam1* mutation. Panel C shows segregation of the deletion in progeny from a cross between ‘Anna’ and ‘Reverend Morgan”. The double band indicates the presence of the deletion, as inherited from the heterozygous ‘Anna’ parent. The raw gel image is provided in the supplementary file.

### Sequence Diversity Among ‘Shell of Alabama’ Accessions

The ‘Shell of Alabama’ genotype is well known as a pollinizer for early-season flowering ‘Anna’ and ‘Dorsett Golden’, consistent with the finding of a common mutation in a known regulator of dormancy. Yet observations of ‘Shell of Alabama’ indicate that not all trees of this variety emerge from dormancy in sync with other ULC varieties, typically flowering weeks later. To test the hypothesis that there are at least two different genotypes sharing this variety name, independent ‘Shell of Alabama’ trees were sampled from material obtained from reputable sources across a broad geographical region (Figure 5). Trees were classified as flowering with ‘Anna’ or after (Table II). The trees that flowered early also presented the *dam1* mutations and the co-segregating 36 bp deletion. Late flowering trees did not contain the mutations or the deletion. Genome sequence was compared from five ‘Shell of Alabama’ individual trees to infer relationships. The results show that the early-flowering ‘Shell of Alabama’ trees (Shell 4 and Shell 7) are near siblings and highly related. The late-flowering accessions (Shell1, Shell 2, Shell 5) also appear to be siblings. Kinship analysis shows that the early-flowering trees are foundational genotypes relative to the later-flowering individuals (Table III).

**Figure 5.**
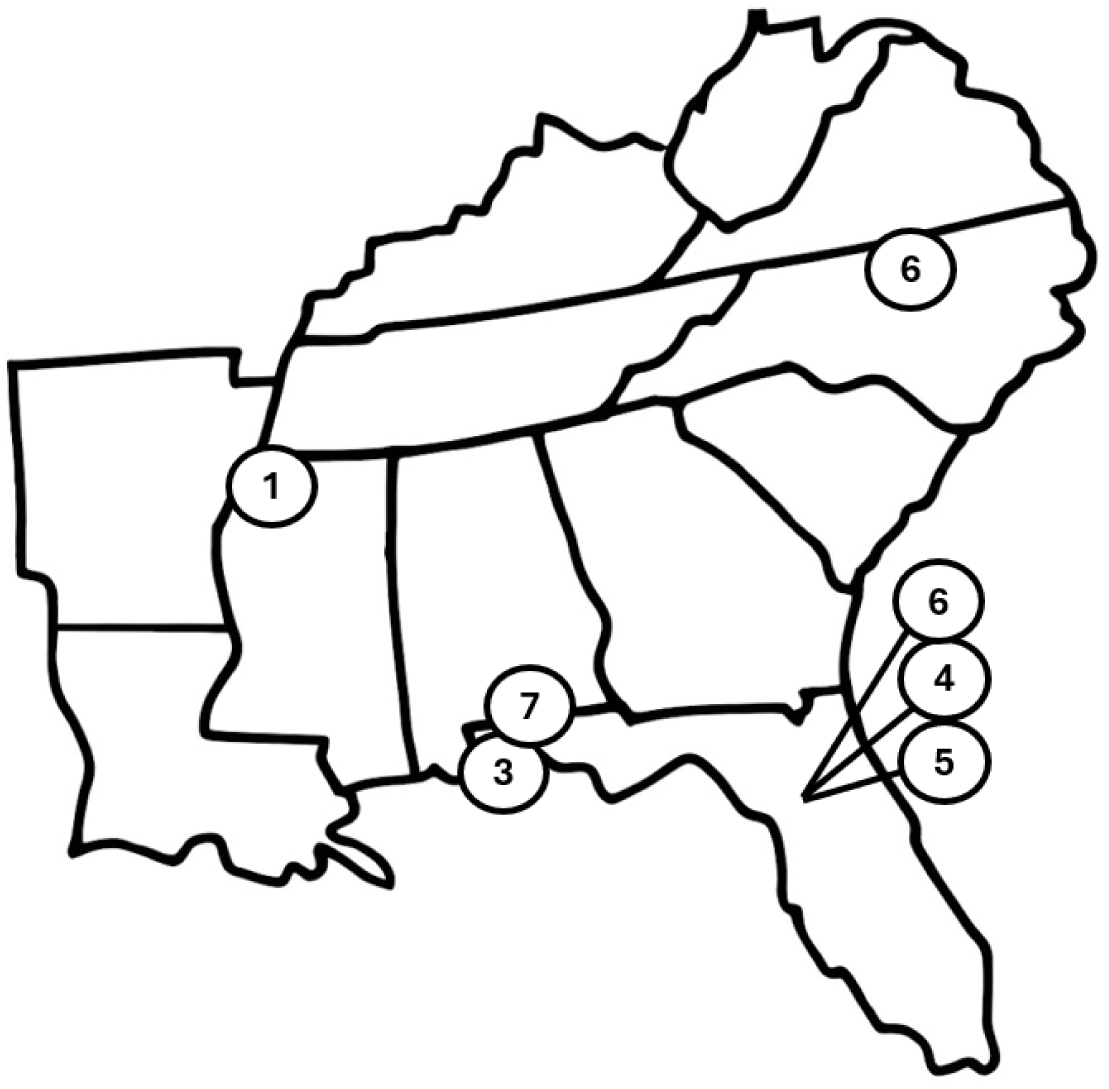
The geographical origins of “Shell of Alabama” samples.

**Figure 5.**
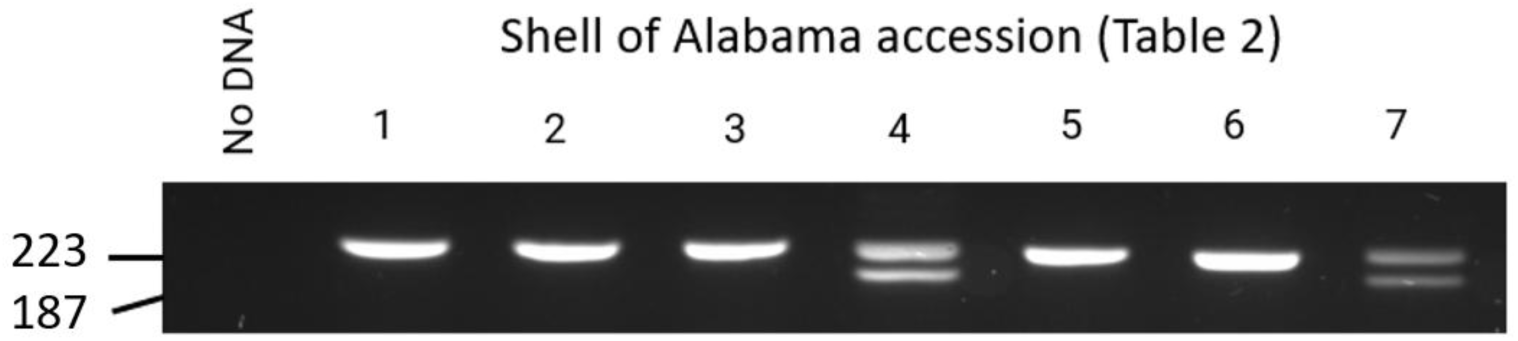
The deletion that co-segregates with the ultra-low-chill phenotype is detected in two of the ‘Shell of Alabama’ accessions tested. The 223 bp PCR product represents the wild-type allele, the 187 bp product segregates with early emergence from dormancy.

**Table II.** The ‘Shell of Alabama’ materials used in this analysis.

| Samples | Tree Location | Early/late | 36 bp<br>Deletion | DAM1<br>Status | Genome<br>Sequenced |
| --- | --- | --- | --- | --- | --- |
| Shell 1 | Clearwater, MS | Late | Absent | Normal | Yes |
| Shell 2 | Archer, FL | Late | Absent | Normal | Yes |
| Shell 3 | Pace, FL | Late | Absent | Normal | No |
| Shell 4 | Archer, FL | Early | Present | Mutant | Yes |
| Shell 5 | Archer, FL | Late | Absent | Normal | Yes |
| Shell 6 | Reidsville, NC | Late | Absent | Normal | No |
| Shell 7 | Flomaton, AL | Early | Present | Mutant | Yes |

**Table III.** Relationships between ‘Shell of Alabama’ based on genomic sequence (individuals 1, 2, 4, 5, 7 in Table II)

| Sample 1 | Sample 2 | Z0 | Z1 | Z2 | PI_HAT | Relationship |
| --- | --- | --- | --- | --- | --- | --- |
| Shell 4 | Shell 7 | 0 | 0.8 | 0.2278 | 0.6139 | Full siblings |
| Shell 4 | Shell 1 | 0 | 0.9 | 0.0989 | 0.5494 | Parent-offspring |
| Shell 4 | Shell 5 | 0 | 1.0 | 0 | 0.5 | Parent-offspring |
| Shell 4 | Shell 2 | 0 | 0.9 | 0.0972 | 0.5486 | Parent-offspring |
| Shell 1 | Shell 5 | 0 | 0.8 | 0.1871 | 0.5935 | Parent-offspring or full siblings |
| Shell 1 | Shell 7 | 0 | 0.9 | 0.1463 | 0.5732 | Parent-offspring |
| Shell 2 | Shell 5 | 0 | 0.8 | 0.1964 | 0.5982 | Full siblings |
| Shell 2 | Shell 7 | 0 | 0.8 | 0.1507 | 0.5753 | Parent-offspring |
| Shell 5 | Shell 7 | 0 | 1.0 | 0 | 0.5 | Parent-offspring |

**Table IV.**
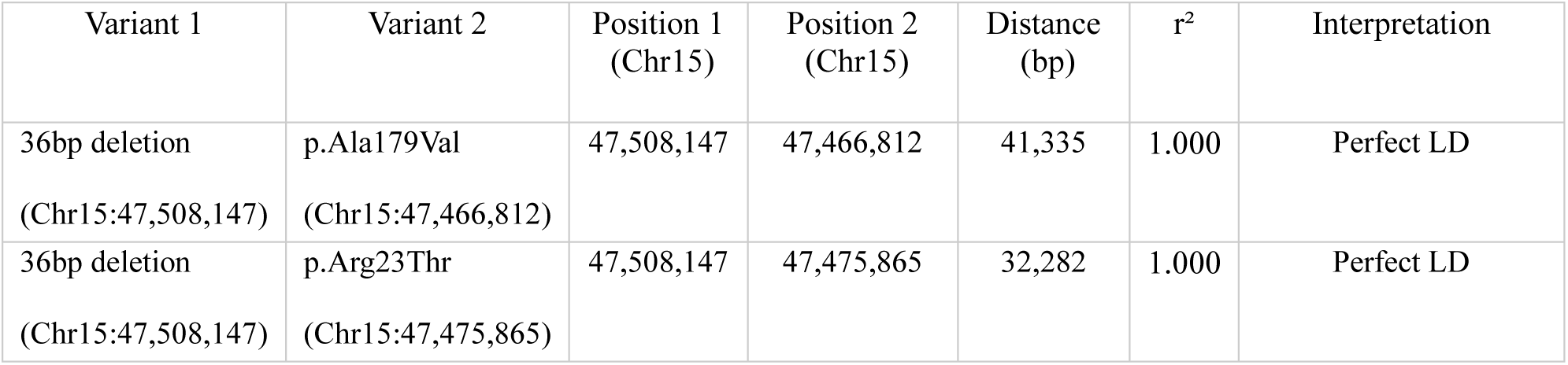
Pairwise linkage disequilibrium (r²) among *DAM1* low-chill variants and the 36 bp lineage marker.

| Variant 1 | Variant 2 | Position 1<br>(Chr15) | Position 2<br>(Chr15) | Distance<br>(bp) | $r^2$ | Interpretation |
| --- | --- | --- | --- | --- | --- | --- |
| 36bp deletion<br>(Chr15:47,508,147) | p.Ala179Val<br>(Chr15:47,466,812) | 47,508,147 | 47,466,812 | 41,335 | 1.000 | Perfect LD |
| 36bp deletion<br>(Chr15:47,508,147) | p.Arg23Thr<br>(Chr15:47,475,865) | 47,508,147 | 47,475,865 | 32,282 | 1.000 | Perfect LD |

## Discussion

The present work centers primarily on three well-established, ultra-low-chill varieties that are commonly used as pollinizers of each other because of their simultaneous early flowering-- ‘Anna’ from Israel, ‘Dorsett Golden’ from the Bahamas, and ‘Shell of Alabama’ from southern Alabama USA. This triad, in conjunction with ‘Tropic Sweet’ (a descendant of ‘Anna’) and ‘Ein Shemer’, and several other genotypes, represent the most commonly planted varieties in tropical and sub-tropical regions with low chill hour environments. However, phenological observations suggested that ‘Anna’ and ‘Dorsett Golden’ are more similar than the popular narrative suggests, likely arising from a common lineage in Israeli breeding efforts (Miller and Sherman, 1980). Molecular analyses identified common variant patterns and genetic relationships suggesting that ‘Dorsett Golden’ is the offspring of ‘Anna’ and another unknown parent (Muranty et al, 2020). An analysis of 73 low-chill historical varieties in the Israeli breeding program identified a shared 190 kb region on chromosome 9 which supposedly contribute to the low-dormancy phenotype (Hauagge & Cummins, 2000; Trainin et al, 2016; van Dyk et al, 2010).

Our first efforts to clarify the relationship utilized an established set of primers to examine microsatellite, or ‘simple sequence repeat’ (SSR) variation between common low-chill apple varieties. The findings of SSR analysis support the hypothesis of Miller and Sherman (1980), that ‘Dorsett Golden’ is highly related to ‘Anna’. It is less consistent with the popular story that low-chill apple originated from a lone seedling of ‘Golden Delicious’ apple cultivar in the Bahamas. Patterns of relatedness were not completely unexpected because both ‘Dorsett Golden’ and ‘Anna’ allegedly share ‘Golden Delicious’ as a common parent.

But to our surprise, these varieties also showed remarkable similarities to ‘Shell of Alabama’, an apple variety presumed to be of completely separate lineage, as it preceded the Israeli ULC varieties by 80-years and originated and was cultivated in the southern United States. Other southern USA accessions (including but not limited to ‘Elsa Sweet’ and ‘Joy’s Apple’) also demonstrated a common SSR-based pattern, indicating a number of regional selections likely originated from a similar genetic base. Common flowering time and observational data of interfertility also support this hypothesis. In contrast, other varieties described as having a higher chill requirement exhibited clearly different microsatellite patterns compared to ULC varieties. This was somewhat unexpected as the Israeli varieties and other lower-chill varieties (e.g. Gala, Cripp’s Pink) also share ‘Golden Delicious’ as a parent. Overall, SSR analysis was able to delineate the ULC genotypes apart from other higher-chill varieties, suggesting common lineage among the ULC varieties, despite their presumably disparate origins in time and location.

Genome sequencing provided a higher resolution interrogation of the ULC genotype relatedness. Analysis of global SNP variation reinforced the close relationship between ‘Anna’ and ‘Shell of Alabama’, with slightly less similarity to ‘Dorsett Golden’. This analysis aligns with the SSR findings and close relationship within ULC genotypes, reaffirming a common lineage. The degree of similarity suggests that these may be vegetatively propagated versions of each other, or possibly self-fertilization, yet they do maintain many SNP differences. These accessions also show differences in fruit phenotypes that could not likely be explained solely by vegetative propagation variation of a sport.

Although ‘Anna’ and ‘Dorsett Golden’ are recognized as offspring of ‘Golden Delicious’, there was no clear evidence of ‘Golden Delicious’ as the parent of ‘Dorsett Golden’. This discrepancy is particularly striking since ‘Dorsett Golden’ has been popularly described as a seedling resulting from self-pollination of ‘Golden Delicious’, isolated because if flowered in the Bahamas. The lack of genome-level confirmation for the presumed parent of ‘Dorsett Golden’ underscores the need for further investigation.

When the genomic DNA sequence of ULC varieties was compared to non-ULC genotypes, and variations were queried in candidate dormancy and flowering genes, a likely causal gene emerged. SNPs were detected that affected the amino acid sequence of the *DAM1* gene, a member of the *DORMANCY ASSOCIATED MADS-BOX* multigene family of transcription factors with well described roles in dormancy control. Much of what has been learned about maintenance of dormancy, especially in rosaceous crops, comes from peach (*Prunus persica*) where the *evergrowing* mutant revealed the genetic basis of vegetative growth repression (Bielenberg et al, 2004). The *evergrowing* mutant continued vegetative growth into short days and falling temperatures without entering into dormancy or setting terminal buds (Jiménez et al, 2009; Rodríguez-A et al, 1994). Genetic analysis identified a 132 kb deletion a cluster of six *DAM* genes, noted as *ppDAM1-ppDAM6* (Bielenberg et al., 2004). They are part of the *SHORT VEGETATIVE PHASE* (*SVP*) family of genes, a set of genes regulating transition to flowering Arabidopsis, that have roles in the regulation of dormancy in deciduous trees (Yamane *et al*, 2021).

In peach, *ppDAM5* and *ppDAM6* serve as repressors of genes associated with growth, and they are highly expressed (especially *ppDAM6*) during deep dormancy (Zhao et al, 2023). Loss-of-function mutation of these genes leads to the *evergrowing* phenotype. Apple also maintains *DAM* gene orthologs with similar function. They work in tandem with a series of DNA binding activities to modulate a predictable suite of transcripts with roles in growth and response to environmental stress (Falavigna et al, 2019; Falavigna et al, 2021; Miotto et al, 2019). The *DAM* genes are encoded by small multigene clusters present on separate chromosomes. One locus on Chromosome 9 has been implicated in dormancy control (Trainin et al. 2016). However, expression data indicate that this paralog is not expressed under dormancy conditions (Moser et al, 2020) and does not possess the domains found in *DAM1* on Chromosome 15. An accounting of transcripts with differential accumulation in *MdDAM1* overexpression and RNAi transgenics shows that Chromosome 15 paralog (MD15G1384500) has abundant transcript accumulation during dormancy (Moser et al, 2020). The same chromosome 15 paralog is the gene featuring shared loss-of-function *dam1* mutation in ULC varieties.

The mutation of *DAM1* in three ULC varieties, not found in other higher-chill-requirement varieties, provides reasonable evidence for *MdDAM1* on Chromosome 15 as a gene with a central role in maintenance of endodormancy. These findings align with the loss-and gain-of-function experiments performed by Moser et al. (2020) that recognize *MdDAM1* as the causal repressor of seasonal growth using functional transgenic experiments. These results are consistent with their role in other rosaceous crops, such as peach (Jiménez et al., 2009), Japanese apricot (*Prunus mume*; Hsiang et al, 2024; Yamane, 2014), European plum (*Prunus domestica*; Quesada-Traver et al, 2020), and Almond (*Prunus dulcis*; Prudencio et al, 2018) among others. The present work identifies the precise genetic lesions that are predicted to cause loss of function in *MdDAM1*, namely an Arg23Thr mutation that is predicted to reduce DNA binding in the MADS domain, as well as an Ala179Val mutation that is predicted to reduce interactions with co-activators. This hypothesis is compelling because it identifies a common functional root in three genotypes previously thought to have arisen independently across different times and locations. Future efforts will focus on creating the ULC *dam1* mutation in disease resistant or long-storage apple cultivars using gene editing, potentially adapting them to low-chill environments.

The key finding is the common mutations that occur in genotypes thought to be remarkably independent, suggesting a common origin of the ultra-low-chill trait. While the original hypothesis of this work was that the different varieties would possess contrasting mechanisms to limit dormancy, genomic sequence indicates a common origin. A molecular marker was identified that co-segregates with the *dam1* mutation (Figure 4B and 4C). The heterozygous indel is detected in the early-flowering ‘Shell of Alabama’ and Israeli accessions ‘Anna’ and ‘Ein Shemer’, as well as ‘Dorsett Golden’. It is interesting that the latter three share ‘Golden Delicious’ as a paternal parent, yet ‘Golden Delicious’ does not contain the *dam1* mutation or the associated marker (not shown). Pedigrees of ‘Anna’ and ‘Ein Shemer’ indicate different maternal parentage, ‘Red Hadassiya’ and ‘Zabadidni’, respectively. ‘Dorsett Golden’ is alleged to be a seedling of ‘Golden Delicious’. While all these genotypes share a common trait, and a common parent, there is no ‘Golden Delicious’ genotype in publicly available data that contains the mutant *dam1* alleles or the associated deletion, leaving the pedigree potentially in need of revision. It is notable that the ‘Vered’ genotype, also from an Israeli breeding program, does not contain the marker or the mutation, yet emerges from dormancy in sync with ‘Anna’, ‘Ein Shemer’, ‘Dorsett Golden’ and early ‘Shell of Alabama’, indicating that there are additional mechanisms that control emergence from endodormancy, or possibly other *dam1* allele-containing genotypes isolated independently.

In terms of commercial influence of the mutation, a significant acreage of ‘Dorsett Golden’ apples was planted in Florida in the 1970’s only to reach the conclusion that the crop had limited commercial potential in sub-tropical regions (Andersen et al, 2022; Miller & Sherman, 1980). However, half a century later there is a newfound interest in low-chill apples, with successful operations flourishing throughout the tropics and sub-tropics. This has enabled the introduction of new disease-tolerant, vigorous rootstocks, the discovery of southern-USA-grown local varieties that flower and produce early, and the development of new varieties for low-chill cultivation. The ULC trait is known to be dominant in segregating populations (Hauagge & Cummins, 2000), which may accelerate variety improvement. Coupled to a robust molecular marker, improved rootstock vigor, and management techniques that may accelerate flowering, it invites the possibility to rapidly develop new varieties to help farmers diversify or even anchor small commercial operations. In addition to new varieties for the tropics and sub-tropics, such analyses may inform development of new varieties for temperate regions as global mean temperatures increase.

The variation identified within ‘Shell of Alabama’ individuals arose from observations of in-orchard trees that flowered at distinct times relative to ‘Anna’ and ‘Dorsett Golden’. Five independent genomes were sequenced and compared, and markers were evaluated for six trees total. Two genotypes show the high similarity to ‘Anna’ and ‘Dorsett’, and kinship analysis suggests that the early ‘Shell of Alabama’ precedes these other varieties. The later flowering ‘Shell of Alabama’ appears to be an offspring of the ULC ‘Shell of Alabama’, showing a sibling relationship with ‘Anna’ and ‘Dorsett Golden’. These relationships suggest that the late ‘Shell of Alabama’ genotype arose as a seedling of the ULC ‘Shell of Alabama’, which is heterozygous for the *dam1* mutation. It is tempting to speculate that the high-chill ‘Shell of Alabama’ was selected from seedlings arising from self-pollination, somewhere where late spring freezes would limit successful flowering and fruiting from a ULC genotype. The variations observed between apparent siblings are not surprising, as these are likely clonally propagated over time and accumulated SNP variations (Li, 2016; Vondras *et al*, 2019).

The most foundational accessions are ULC, possess the *dam1* mutation, and flower in unison with ‘Anna’ and ‘Dorsett Golden’. One of these is located in Flomaton, AL, in the region of central Alabama near the Florida state line where Greene Shell’s orchards produced commercial apples in the late 1800’s, suggesting it is the *bona fide* genotype, bolstering the relationships inferred from genome sequence comparisons. Future studies will analyze additional genomes from the region, testing for the presence of the *dam1* mutation in the oldest orchards and residential trees.

The limitations of this study are in the paucity of ULC genotypes in central repositories, and dependence on decades of chain-of-custody to maintain variety integrity. Additional work will examine the correlation of the *dam1* mutations and flowering time in formal trials, as well as attempt to install the *dam1* mutations in highly disease resistant germplasm. Genomes of other ULC genotypes will be sequenced and analyzed to further illuminate the relationships between ULC varieties.

In conclusion, analysis of the mechanism beneath the ULC phenotype in three popular sub-tropical/tropical apple varieties led to several important findings. A predicted genetic relationship between ‘Anna’ and ‘Dorsett Golden’ is confirmed, and both are now linked to an early-flowering ‘Shell of Alabama’ as a near-relative. Identification of the precise genetic lesion likely underlying the ULC response provides traceable molecular marker for introgression of the dominant apple *dam1* mutations into future low-chill breeding efforts. The variation in ‘Shell of Alabama’ accessions suggests loose chain-of-custody of the original genetics, and possibly propagation by seed. The curious absence of the trait and associated marker from ‘Golden Delicious’ sequence, despite being the presumed parents of ‘Anna’, ‘Ein Shemer’ and ‘Dorsett Golden’, also merits further inquiry.

## Acknowledgements

The authors thank Larry Stephenon (Southern Cultured Nursery, Coldwater, MS), Randall White (Flomaton, AL), Ron Fairbanks (Pace, FL) and Barry Overton (Archer, FL; with a tree originating from North Carolina) for their contributions of known ‘Shell of Alabama’ material and their keen observations.

## Data Availability

All genome sequence data is present in the Short Read Archive under the following accession numbers: (we will add these upon acceptance, with appropriate embargo)

