## Supplementary File for "Identical Dormancy Gene Mutations Reveal Unanticipated Relatedness Among Low-Chill Apples"

**Extraction of nuclei**

The fresh leaves were used for the nuclei extraction with modified method as described by (Li et al., 2020) . All the buffers/solutions were prepared as described by (Li et al., 2020) . The 5 g leaves were ground with the mortar and pestle using liquid nitrogen and the fine powder was transferred to 500 ml flask containing 30 ml cold, fresh prepared, Nucleic Isolation Buffer (NIB). The 0.5% 2-Mercaptoethanol (BME) was added in NIB buffer before use. The flask containing a mixture of ground sample and NIB was gently shaken at 100 rpm for 10 minutes on ice. For filtration, the mixture was passed through two layers of Cheesecloth and two layers of Mira-cloth with the help of a 15 cm diameter funnel. The Mira-cloth was touching the flask while Cheesecloth was kept on top of it as the first layer of filtration. The filtrate was allocated into two 50 ml conical tubes and centrifuged at 2500 x g at 4 °C for 12 minutes. The pellet was resuspended with the help of a small paint brush and 3 ml cold NIB was added in each tube. Again, centrifuged at 2500 x g at 4 °C for 6 minutes. The pellet was resuspended with a small paint brush by adding 0.5 ml NIB and the suspension was transferred into 2ml Eppendorf. The suspension was centrifuged at 3000 x g for 15 minutes at 4 °C. The washing with NIB with 0.5 ml NIB was done 3-5 times until clear white suspension was not achieved. After attaining white suspension, the pellet was suspended with 0.2 ml NIB and suspension of both Eppendorf was transferred into one new clean 15 ml conical tube. The 4 ml pre-heated (65 °C) 2 x CTAB buffer with 0.5% BME was added, mixed well gently and cooled down to room temperature. After that, equal volume of chloroform was added, mixed well gently, and centrifuged 2500×g for 15 minutes at room temperature.

The supernatant was transferred to a new clean 15 ml conical tube 0.5 ml 10% CTAB buffer was added. The mixture was mixed well and incubated at 65 °C for 5 minutes. After cooling down to room temperature, the equal volume of chloroform was added and gently mixed by inversion. The mixture was centrifuged at 2500×g for 15 minutes at room temperature and the supernatant was transferred into a new clean 15 ml conical tube. The genomic DNA was precipitated by adding the equal volume of 1 x CTAB buffer and inverting till white fluffy DNA is visible. The centrifugation at 2500×g for 1 minute was done to precipitate the DNA and pellet was resuspended in 200 µl of High-salt T.E solution. The suspension was transferred into 2ml tube 700 µl ethanol was added, incubated at room temperature for 3 minutes. Again, the genomic DNA was precipitated by centrifugation at 3000×g at room temperature for 3 minutes. The pellet was washed twice by adding 700 µl and spinning 3000×g at room temperature for 3 minutes. After airdrying pellet for 20-40 minutes, 100 µl 0.1 x T.E buffer was added.

**DNA Extraction**

1. The 3mg leave sample was ground using liquid nitrogen and 800 ul of CTAB buffer (pre-heated) was added.
2. The mixture was incubated at 58 °C for 2 hours.
3. After that 800 ul of Chloroform: isoamyl alcohol (24:10 ratio was added and mixed well by shaking.
4. The mixture was centrifuged at 14,000 rpm for 15 minutes.
5. The white upper aqueous layer was transferred to new tube.
6. The 0.6 volume of isopropanol was added and incubated overnight for DNA precipitation.
7. The DNA was washed using 70% ethanol first and then with 95%.
8. The samples were air dried and DNA pellets were suspended in T.E buffer for downstream analysis.

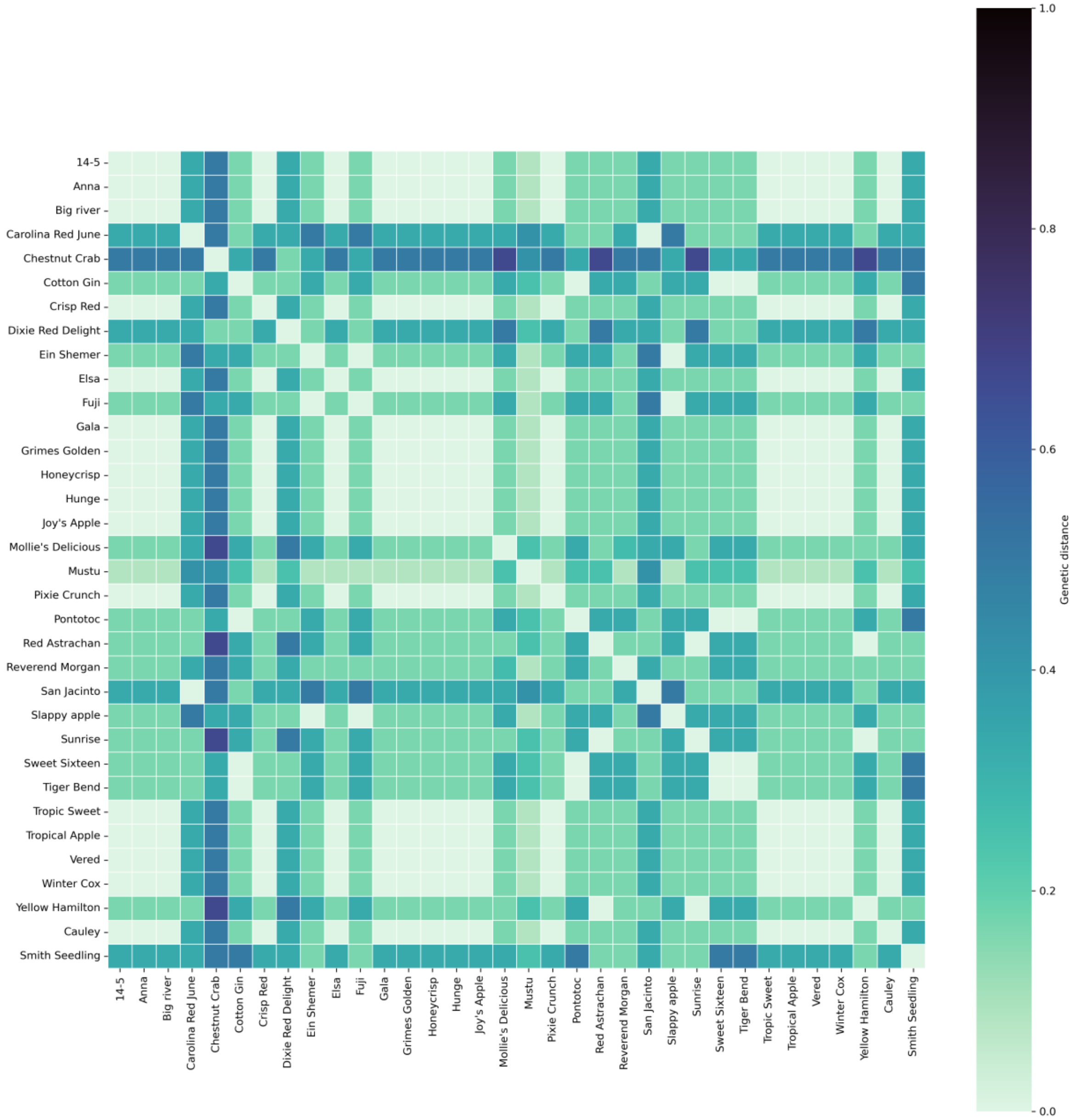


Supplimentary Figure No.1. Shows the relatedness among Southeren apple cultivars based on allelic similarity. The colored scale bar (0-1) shows higher distance among the cultivars.

Supplementary Table No. 1. Fluorescence tagged primers used for SSR analysis.

| No. | SSR marker | Primer sequence | Product length | Chromosome locus |
|---|---|---|---|---|
| 1 | CH02c06 | F: AGTTTCGTAAGAGAACCTTGATCTC R: ATCCACTTACTAAGAACTACCGTTG | 400-450 | Chr: 2 |
| 2 | GD12 | F: GAATGTGAGGCGTTCCTGAG R: GAATGCGAGGCCTTCCTGAG | 300-350 | Chr: 3 |
| 3 | CH04e05 | F: GAGAAGGCTAACAGAAATGTGG R: GCCTTTGTAATCATGGCTCC | 200-250 | Chr: 7 |
| 4 | CH01h10 | F: GCAAAGATAGGTAGATATATGCCA R: AAGGAGGGATTGTTTGTGCA | 100-150 | Chr: 8 |
| 5 | CH02c11 | F: CTGAGGTATTATTTTGTTTCTGCG R: GAAACACATTTATAGAAAAAGGAGC | 100-150 | Chr: 10 |

Supplementary Table No.2. Plink based genetic relationship among apple cultivars.

| IID1 | IID2 | Z0 | Z1 | Z2 | PI_HAT | DST |
|---|---|---|---|---|---|---|
| Anna | Dorsett Golden | 0 | 0.9 | 0.1 | 0.57 | 0.81 |
| Anna | Shell | 0 | 0.0 | 1.0 | 0.98 | 0.99 |
| Anna | Gala | 0 | 1.0 | 0.0 | 0.50 | 0.75 |
| Anna | Red Delicious | 0 | 0.6 | 0.4 | 0.71 | 0.87 |
| Anna | Antonovka | 0 | 0.8 | 0.2 | 0.60 | 0.82 |
| Anna | Honeycrisp | 0 | 1.0 | 0.0 | 0.51 | 0.78 |
| Dorsett Golden | Shell | 0 | 0.9 | 0.1 | 0.57 | 0.81 |
| Dorsett Golden | Gala | 0 | 0.8 | 0.2 | 0.58 | 0.81 |
| Dorsett Golden | Red Delicious | 0 | 1.0 | 0.0 | 0.50 | 0.78 |
| Dorsett Golden | Honeycrisp | 0 | 0.8 | 0.2 | 0.60 | 0.82 |
| Gala | Red Delicious | 0 | 1.0 | 0.0 | 0.50 | 0.77 |
| Gala | Honeycrisp | 0 | 0.9 | 0.1 | 0.55 | 0.80 |
| Antonovka | Honeycrisp | 0 | 1.0 | 0.0 | 0.52 | 0.79 |

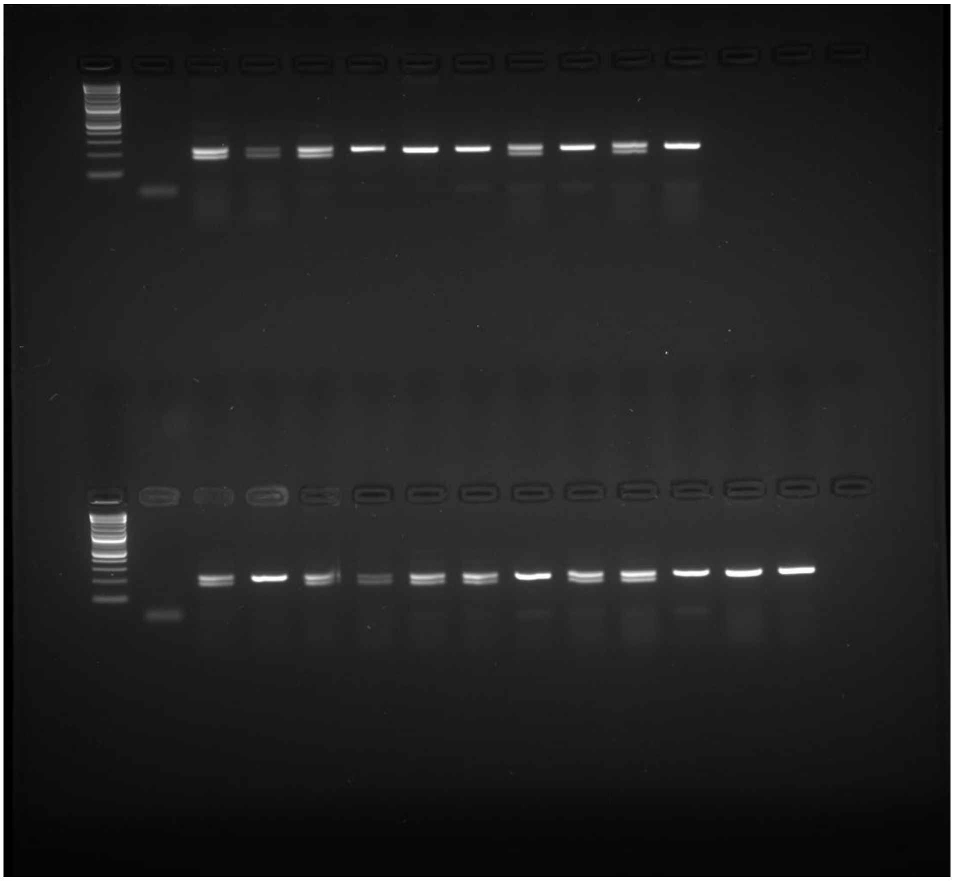

Supplementary Figure No. 2. Raw gel image used for figure 4b and 4c in the main text.

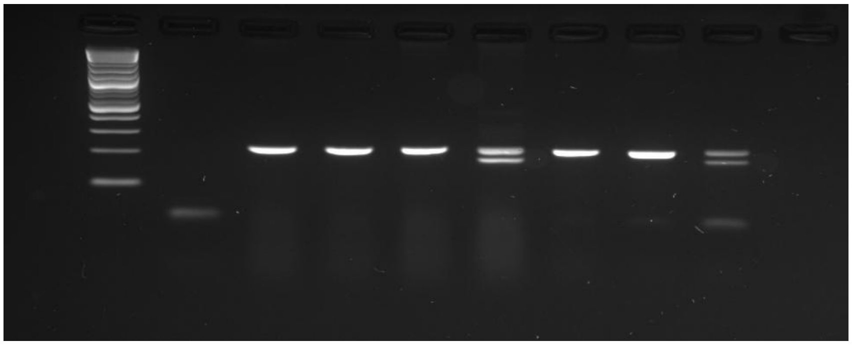

Supplementary Figure No. 3. Raw gel image used for figure 5 in the main text.
